# Quantifying Rift Valley fever virus transmission efficiency in a lamb-mosquito-lamb model

**DOI:** 10.1101/2023.04.25.538218

**Authors:** Gebbiena M. Bron, Paul J. Wichgers Schreur, Mart C.M. de Jong, Lucien van Keulen, Rianka P.M. Vloet, Constantianus J.M. Koenraadt, Jeroen Kortekaas, Quirine A. ten Bosch

## Abstract

Rift Valley fever virus (RVFV) is a (re)emerging mosquito-borne pathogen impacting human and animal health. How RVFV spreads through a population depends on population-level interactions between hosts and vectors (e.g., vector-to-host ratio and biting preference) and also on potential differences in individual following virus exposure (e.g., transmission efficiencies from host to vector and vice versa). Here, we estimated the probability for RVFV to transmit to naive animals by experimentally exposing lambs to a bite of an infectious mosquito (the transmission efficiency) and assessed if and how RVFV infection subsequently developed in the exposed animal. *Aedes aegypti* mosquitoes, previously infected via feeding on a viremic lamb, were used to expose naive lambs to the virus. Lambs were either exposed to 1-3 (low exposure) or 7-9 (high exposure) infectious mosquitoes. All lambs in the high exposure group became viremic and showed characteristic signs of Rift Valley fever within 2-4 days post exposure. In contrast, 3 out of 12 lambs in the low exposure group developed viremia and disease, with similar peaks in viremia as the high exposure group but with some heterogeneity in the onset of viremia. These results suggest that the likelihood for successful infection of a ruminant host is affected by the number of infectious mosquitoes biting, but also highlights that a lamb can be infected by a single mosquito. The per bite mosquito-to-host transmission efficiency was estimated at 28% (95% confidence interval: 15 - 47%). We subsequently combined this transmission efficiency with estimates for mosquito life traits into a Ross-McDonald mathematical model to illustrate scenarios under which major RVFV outbreaks could occur in naïve populations (i.e., R_0_ >1). The model revealed that for efficient RVFV transmission relatively high vector-to-host ratios as well as strong feeding preference for competent hosts are required. Altogether, this study highlights the importance of experiments that mimic natural exposure to RVFV. The experiments facilitate a better understanding of the natural progression of disease and a direct way to obtain epidemiological parameters for mathematical models.

## Introduction

Rift Valley fever virus (RVFV) is a mosquito-borne, zoonotic virus within the *Phenuiviridae* family, order of *Bunyavirales*, that mostly affects ruminants (Daubney et al., 1931). In ruminants, the virus may cause abortion storms and juvenile mortality, with sheep being most affected (Findlay, 1932; Ikegami and Makino, 2011). In humans, Rift Valley fever (RVF) occurs incidentally. The disease is often sub-clinical but can be fatal. Humans become infected by RVFV through contact with infectious tissues, e.g., at slaughter or during abortion by contact with placenta and fluids, or after being bitten by infectious mosquitoes. Over 40 species of mosquito have been found competent to transmit RVFV under laboratory conditions. This contributes to the potentially wide geographic range of the virus. Due to the impact of RVFV outbreaks on livestock and trade, the virus has a major impact on a region’s economy and livelihoods of millions of people (Tempia and Abdi, 2006; Chengula et al., 2013). Due to its potential to cause a public health emergency in absence of efficacious drugs and (human) vaccines, the World Health Organization identified RVF as a disease in urgent need for accelerated research and vaccine development (Mehand et al., 2018). In addition, its recent emergence into new regions, such as the Arabian Peninsula, urges the need to better understand the potential of RVFV to spread beyond its original habitat (Bron et al., 2021).

The ability of RVFV to cause outbreaks and establish in new places depends on many factors, including the nature and frequency of interactions between vectors and vertebrate hosts (Figure 1). The probability that an interaction (through a mosquito bite) between infected and susceptible hosts and vectors results in onward transmission of the pathogen is captured by a metric called transmission efficiency. For RVFV, various experimental studies have contributed to an increased understanding of the ability of vertebrate hosts and mosquito vectors to transmit the virus. Yet, these studies are less suited for informing the efficiency of pathogen transmission.

**Figure 1.**
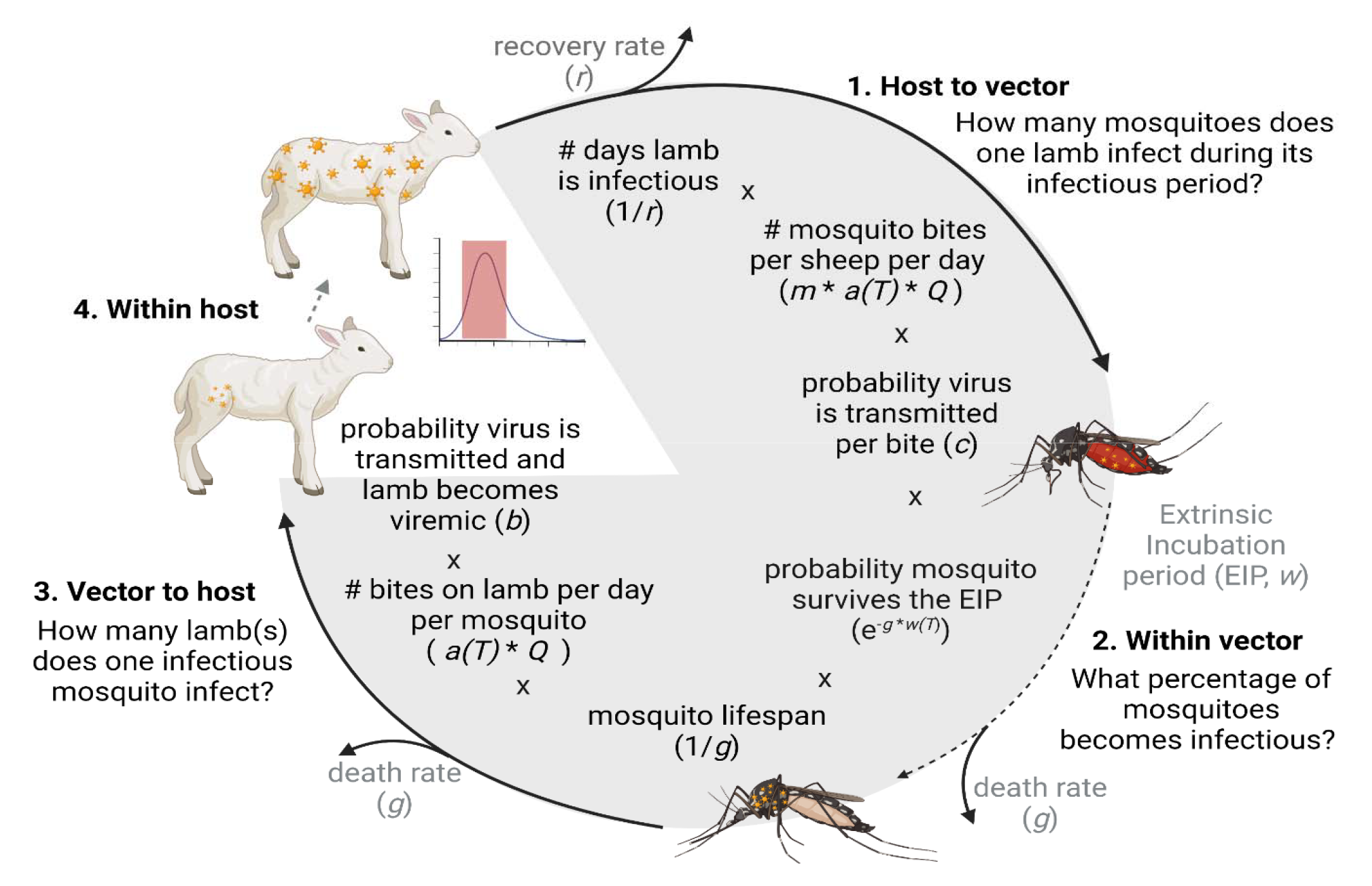
A model of the RVFV transmission cycle and the components of the basic reproduction number (R_0_). How many new viremic lamb(s) originate from one viremic lamb? Here we follow the virus from host to vector (1), within the vector (2), from vector to host (3) and within the host (4). First, we calculate how many mosquitoes will feed on a viremic lamb and how many of those will become infected (1). Next we estimate how many mosquitoes survive to become infectious (2). We then estimated, for those surviving mosquitoes, how many sheep they will bite over the remainder of their live, whereby we assume that mosquitoes do not clear the infection. Multiplying this number with the probability of successful transmission from mosquito to sheep (*b*) gives the total number of sheep to be infected by a single infected mosquito. This number is also referred to as the type reproduction number for mosquitoes (3. Vector to host). In our experiment we estimated *c* (the probability that virus is transmitted from a viremic sheep to a mosquito), *b* (the probability a sheep is successfully infected and becomes viremic after an infectious mosquito bite) and we assessed the within host dynamics of RVFV after low and high mosquito exposure. The other parameters were extracted from literature or set to scenario specific values. Note that while the number of bites a sheep receives depends on the vector to host ratio (*m*), the number of bites a mosquito takes on sheep does not. This reflects the assumption that, provided that the number of hosts is not limiting the mosquito’s feeding behavior, the number of bites a mosquito takes does not increase when more hosts are available. In contrast, an infected sheep will receive more bites if there are fewer other hosts that mosquitoes can distribute their bites over. *m* = vector to host ratio; *a(T)* temperature dependent biting rate including adjustment for multiple biting behavior (number of bites per mosquito per day); *Q* the proportion of bites on sheep (competent host) versus non-competent hosts, this parameter is affected by host availability and mosquito host preference.

Competence studies often provide an incomplete understanding of the efficiency of pathogen transmission, due to, for example, the artificial nature of RVFV exposure or indirect transmission metrics used (e.g., virus in saliva). Furthermore, these studies did illustrate that the outcomes of infection is not only dependent on the vertebrate host species used (Smithburn et al., 1949; Turell et al., 2008), but the outcome is also affected by 1) the mode of RVFV challenge (e.g., needle vs. mosquito bite, mosquito species) (Le Coupanec et al., 2013) 2) the source of the virus, e.g., different cell lines or mammalian hosts (Turell, 1988; Weingartl et al., 2014; Wichgers Schreur et al., 2021), and 3) the dose of the inoculum. The development of a model system that mimics the transmission of RVFV between mosquito vectors and its natural hosts could help resolve such issues.

Studies that investigate the transmission of a virus from a mammalian host to a mosquito vector and back to the mammalian host address all parts of an arbovirus transmission cycle (Figure 1): 1) virus transmission from host-to-vector, 2) vector infection and virus dissemination within the vector, 3) virus transmission from vector-to-host, and 4) infection and disease development in the host. These studies are particularly powerful because they include many of the natural barriers that pathogens encounter in mammalian hosts and mosquito vectors (e.g., physical and immunological barriers). To ensure successful pathogen transmission, experimental challenge studies to assess pathogenesis or vaccine efficacy often expose animals to a relatively high number of mosquitoes (Wichgers Schreur et al., 2021). However, infection through a single or few mosquito bites is likely more relevant to natural exposure than exposure to many mosquitoes, considering relatively low arbovirus infection prevalence.

Moreover, laboratory studies on mosquito-borne pathogen transmission help provide a better understanding of the population-level impact pathogens may have. Translating experimental findings to meaningful epidemiological projections requires a thorough understanding of how vectors and hosts interact with each other in different environments, as well as reliable estimates of 1) individual-level outcomes of infection, and 2) the efficiency of transmission from host to vector and vice-versa. To fill some of these knowledge gaps, we experimentally reproduced and quantified parts of the host-vector-host RVFV transmission cycle in sheep and *Ae. aegypti*. Specifically, we estimate transmission efficiencies from mosquito-to-host and vice versa using biologically relevant experimental exposure models. Next, we describe clinical outcomes and within-host progression of viremia in infected sheep after high and low mosquito exposure. We conclude by using a Ross-McDonald derived model to illustrate how the transmission efficiency estimates map onto invasion risks of RVF in naive populations, across different scenarios of mosquito-host interactions.

## Material & Methods

### Materials

#### Cells and viruses

Culture media and supplements were obtained from Gibco unless indicated otherwise. Baby Hamster Kidney (BHK-21) cells were routinely maintained in Glasgow minimum essential medium (GMEM) supplemented with 4% tryptose phosphate broth, 1% minimum essential medium nonessential amino acids (MEM NEAA), 1% antibiotic/antimycotic (a/a) and 5% fetal bovine serum (FBS), at 37°C with 5% CO_2_.

The challenge virus stock of RVFV strain 35/74, a strain originally isolated from the liver of a sheep that died during a RVFV outbreak in the Free State province of South Africa in 1974 (Barnard, 1979), was obtained by low multiplicity of infection (MOI: 0.005) of BHK-21 cells in the presence of CO_2_-independent medium (CIM, Invitrogen), supplemented with 5% FBS (Bodinco) and 1% Pen/Strep (Invitrogen) (Kortekaas et al., 2011).

For assessment of the presence of RVFV in saliva samples Vero E6 (ATCC, CRL-1586) African green monkey kidney were used. Vero cells were routinely maintained in Earl’s minimum essential medium (MEM) Earl’s supplemented with, 1% L-glutamine, 1% minimum essential medium nonessential amino acids (MEM NEAA), 1% antibiotic/antimycotic (a/a) and 5% fetal bovine serum (FBS), at 37°C with 5% CO2.

#### Mosquitoes and feeding

Rockefeller strain *Ae. aegypti* mosquitoes (Bayer AG, Monheim, Germany) were routinely maintained at the Laboratory of Entomology of Wageningen University and Research (Wageningen, the Netherlands) as described (Visser et al., 2020). Briefly, mosquitoes were kept in Bugdorm-1 rearing cages at a temperature of 27°C with a 12:12 light:dark cycle and a relative humidity of 70%. The mosquitoes were provided with a 6% glucose solution ad libitum.

Mosquitoes were subsequently transported to biosafety level three (BSL-3) facilities of Wageningen Bioveterinary Research (Lelystad, the Netherlands) for the animal experiment.

##### Sheep

Texel-Swifter lambs (*Ovis aries*) were obtained from a conventional sheep farm in the Netherlands. Before inclusion in the study, the general health of lambs was assessed by a veterinarian. Animals were allowed to acclimatize in the BSL-3 facilities for 7 days before the start of the experiment. Food and water were available *ad libitum*.

### Animal experiment

#### Overall study design

A high and low exposure study was conducted using a host-mosquito-host transmission model. Following needle inoculation of 5 lambs with RVFV, mosquitoes *(Aedes aegypti)* were allowed to take a blood meal two days post inoculation, which was previously shown to correspond to peak viremia. Fully engorged mosquitoes were subsequently maintained for 12 days whereafter they were offered a second blood meal this time on naïve lambs. Groups of naive lambs were either exposed to a small number (3; low exposure group) of mosquitoes or a larger group of mosquitoes (28-31 high exposure group). Following mosquito feeding, the progression of infection was assessed in the lambs of the two exposure groups, health status was monitored and viremia levels were determined daily.

#### Ethical approval

The animal experiment was conducted in accordance with European regulations (EU directive 2010/63/EU) and the Dutch Law on Animal Experiments (WoD, ID number BWBR0003081). Permissions were granted by the Dutch Central Authority for Scientific Procedures on Animals (Permit Numbers: AVD4010020185564 and AVD4010020187168). All procedures were approved by the Animal Ethics Committees of Wageningen Research. The following humane endpoints were applied: (1) the animal is recumbent and does not rise even after stimulation, (2) the animal is unable to drink, (3) the animal is lethargic (listless, apathic, non-responsive to stimuli).

#### Obtaining infectious mosquitoes

A group of 10-week-old lambs (N=5) were housed in a BSL-3 animal facility and intravenously inoculated with 10^5^ TCID_50_ of RVFV strain 35/74 at day −14 (Figure 2). Two days post infection (day −12) at expected peak viremia, lambs were sedated by intravenous administration of medetomidine (Sedator, Eurovet Animal Health). When fully sedated, cardboard containers with mosquito netting containing 40–50 naive female *Ae. aegypti* mosquitoes were placed on the shaved inner thigh of each hind leg and mosquitoes were allowed to take a bloodmeal. After approximately 20-30 min of feeding, cardboard containers were removed and animals were euthanized by intravenous injection of an overdose of sodium pentobarbital (Euthasol 20%, AST Pharma). Mosquitoes were transported to a BSL-3 laboratory, where fully engorged mosquitoes were separated from unfed individuals using an automated insect aspirator and pooled in one group. For the next 12 days engorged mosquitoes were maintained in an insect incubator (KBWF 240, Binder) at 28°C, 70% relative humidity and a 16:8 light:dark cycle. The mosquitoes were provided with 6% glucose solution *ad libitum*.

**Figure 2:**
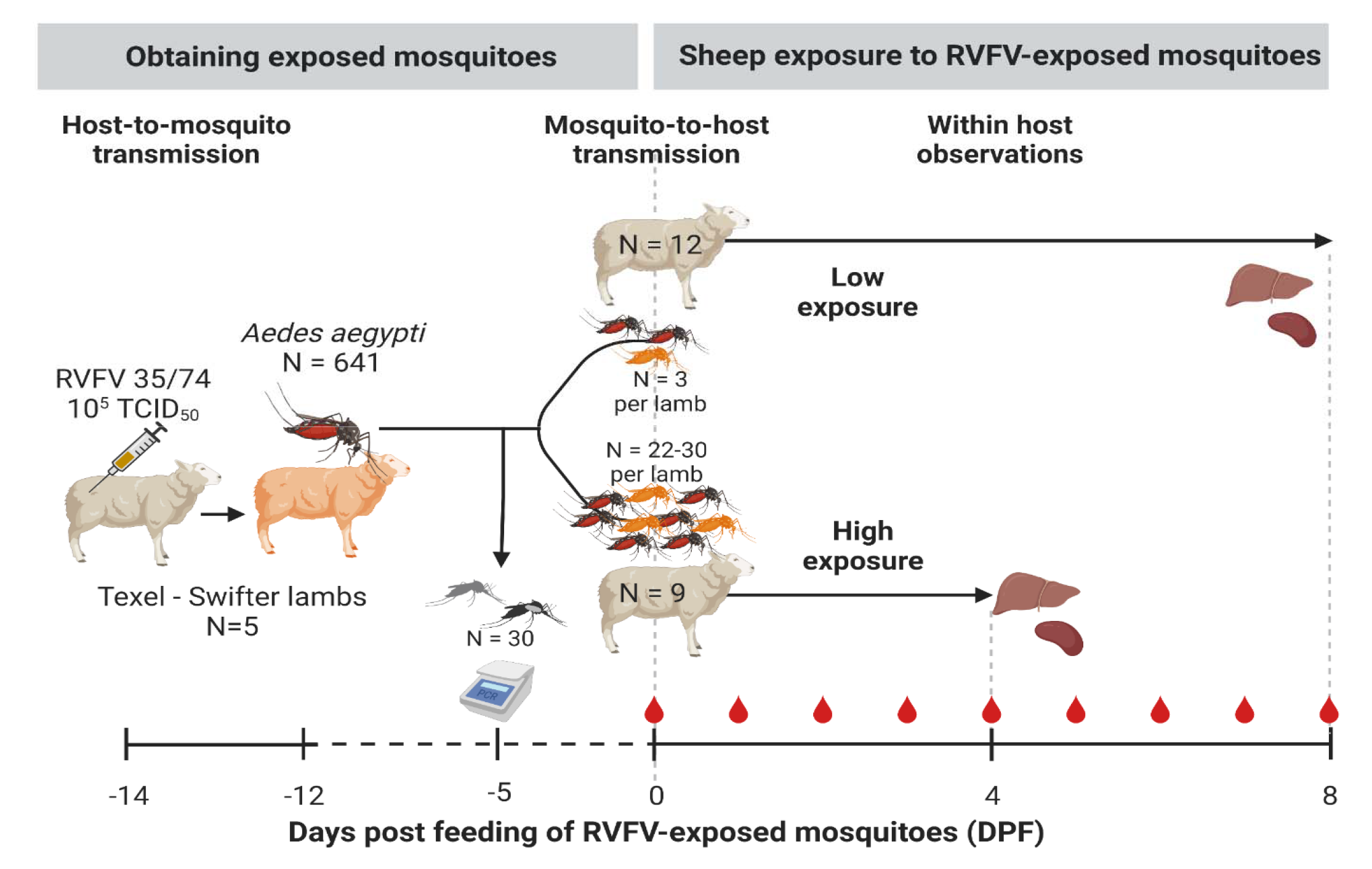
Experimental design. Donor Texel-Swifter lambs were infected by intravenous injection of Rift Valley fever virus (RVFV) two days prior to feeding ∼1000 laboratory reared *Aedes aegypti*. All blood fed mosquitoes (N=641) were subsequently pooled and kept in a climate and light controlled incubator in a BSL3 laboratory. To estimate RVFV infection rates within the population of blood-fed mosquitoes seven days post feeding, 30 mosquitoes were tested for RVFV infection and presence of virus in saliva by RT-qPCR. Twelve days post initial feeding mosquitoes were allowed to take a second blood meal; three (low-exposure group) or 28-31 mosquitoes (high exposure group) were placed in mesh screen containers and allowed to feed on the shaven inner thigh of 47 day-old naive lambs. In the high exposure group 22-30 fed and in the low exposure we ensured three mosquitoes fed. After mosquito exposure, the lambs were monitored 2-3 times daily for abnormalities and rectal temperatures were measured. Blood samples were taken daily (red drop in figure). Animals belonging to the high exposure group were euthanized 4-days post feeding of RVFV-exposed mosquitoes (DPF), to prevent unnecessary discomfort, and low exposed lambs were euthanized at 8 DPF. During necropsy liver and spleen samples were collected for detection of RVFV.

#### Determining the mosquito exposure protocol for low exposure group

To optimize the chance of exposing lamb in the low exposure group to one mosquito with RVFV in saliva (an infectious mosquito), the following protocol was used: At experimental day –5 the RVFV prevalence in a sample group of 30 RVFV-exposed mosquitoes was determined as described in the section ‘sample processing’. The infection prevalence obtained was used to determine the number of mosquitoes allowed to feed per lamb in the low exposure group. After feeding on naïve lamb, the infection status of individual mosquitoes was determined. If no mosquitoes would have tested positive for RVFV, a secondary exposure of animals in the low exposure group was planned two days after the initial exposure date.

#### Exposure of naive lambs to RVFV infected mosquitoes

On day 0 of the experiment, 9 naive 7-week-old lambs were exposed to 28-31 randomly selected mosquitoes that were fed 12 days prior on needle infected lambs (high exposure group, Figure 2). Another 12 lambs were exposed to 3 mosquitoes each (low exposure group). Exposure of the lambs was performed as described in the section ‘Obtaining infectious mosquitoes’.

For the low exposure group, feeding of three mosquitoes was monitored in real-time and when less than three mosquitoes had fed (from the initial container with 3 mosquitoes) additional single mosquitoes were placed on a lamb until 3 mosquitoes had fed on each animal. To assess actual bites per animal for the high exposure group, engorged mosquitoes were separated from unfed mosquitoes and counted within the first day post feeding. Following separation and counting, a subset of 6 mosquitoes per lamb (high exposure group) was randomly selected and subjected to forced salivation and assessment of virus in their bodies. All fed mosquitoes of the low exposure group were tested similarly.

#### Monitoring of mosquito-exposed lambs

Following mosquito exposure, lambs were monitored three times daily for clinical signs and rectal temperatures were measured twice daily. Blood samples were taken on a daily basis to assess viremia (Figure 2). The high exposed lambs were euthanized and necropsied 4 DPF to prevent unnecessary discomfort, anticipating that high exposure would result in severe disease. The low exposed animals were euthanized at 8 DPF. At necropsy, spleen and liver samples were collected.

### Sample processing

#### Lambs

To assess RVFV infection in lambs, RNA was isolated with the NucliSENS easyMAG system according to the manufacturer’s instructions (bioMerieux, France) from 0.5 ml plasma samples. To detect RVFV genetic material RT-qPCR was conducted. Briefly, 5 µl RNA eluate was used in a RVFV RT-qPCR using the LightCycler one-tube RNA Amplification Kit HybProbe (Roche, Almere, The Netherlands) in combination with a LightCycler 480 real-time PCR system (Roche) and the RVS forward primers (AAAGGAACAATGGACTCTGGTCA), the RVAs (CACTTCTTACTACCATGTCCTCCAAT) reverse primer and a FAM-labelled probe RVP (AAAGCTTTGATATCTCTCAGTGCCCCAA). Primers and probes were earlier described by Drosten et al. (Drosten et al., 2002). Virus isolations were performed on RT-qPCR positive samples with a threshold above 10^5^ RNA copies/ml as this was previously shown to be a cut-off point below which no live virus can be isolated.

#### Mosquitoes

To determine if mosquitoes were infected with RVFV, viral RNA was isolated from homogenized mosquitoes and from saliva samples. Briefly, bodies were homogenized in 300 μl Trizol with a pellet pestle (Sigma) and the homogenate was subsequently cleared by slow speed centrifugation. Individual saliva samples (see virus isolation section) were also added to 300 μl Trizol. Total RNA was subsequently isolated using the Direct-zol™ RNA MiniPrep kit (Zymo Research) according to the manufacturer’s instructions. The level of RVFV RNA was subsequently determined as described in section ‘Sample processing; Lambs’.

#### Virus isolation

To determine the presence of infectious RVFV in mosquito saliva, plasma and tissue samples from lambs, virus isolations were performed. To check for positive saliva, mosquitoes were sedated on a semi-permeable CO_2_-pad connected to 100% CO_2_ and wings and legs were removed. Saliva was collected by forced salivation using 20 µl filter tips containing 7 µl of a 1:1 mixture of FBS and 50% sucrose (capillary tube method). After 1–1.5 h, saliva samples were collected and or incubated with Vero-E6 cell monolayers. Cytopathic effect (CPE) was scored 5– 7 days later. To detect RVFV in plasma, serial dilutions of plasma samples were incubated with 20,000 BHK-21 cells/well in 96-wells plates for 1.5 h before medium replacement. Cytopathic effect was evaluated after 5–7 days post infection. Virus titers (TCID_50_/ml) were determined using the Spearman-Kärber algorithm. The limit of detection was 1.55 log10 TCID_50_/ml.

#### Serology

Antibodies against RVFV nucleoprotein (RVFV-NP) were detected by the semi-quantitative ID Screen® Rift Valley Fever Competition Multi-species (IDVet, Grabels, France) enzyme-linked immunosorbent assay (ELISA) weekly. The ELISA was conducted according to the manufacturer’s instructions. Briefly, RVFV-NP was coated in wells, anti-RVFV-NP antibodies in sera were allowed to bind, and anti-NP-peroxidase was used to bind to free RVFV-NP in the well (i.e., RVFV-NP not blocked by anti-NP antibodies). A wash protocol occurred between each step. Finally, substrate (TMB) for bound peroxidase was added, and plates were read out at 450nm. Titers are expressed as percentage inhibition ratio of the optical densities (OD) of the sample and the OD of the negative control (S/N * 100). All values lower than 40% are considered positive, between 40 and 50% are considered doubtful and above 50% are considered negative.

### Data analyses

#### Descriptive statistics

Demographics (age, sex), health status (rectal temperature, survival), and viral infection characteristics (presence of viremia [yes or no], day of onset of viremia, day of peak viremia, and peak viremia level) were summarized. Rectal temperatures (T_r_) from DPF 1 to 4 (last day of study period for the high exposure group) were included to calculate the mean T_r_ and standard deviation per group. A temperature of 40.5°C or higher was considered a fever. To determine the day of fever onset, morning observations were coded as DPF and afternoon observations as DPF + half a day. An animal was considered viremic when the TCID_50_ was above the limit of detection. The probability of developing viremia and mortality were compared between the two exposure groups using Fisher Exact tests (FET). Peak viremia in log_10_ TCID_50_/ml, day of onset of viremia, day of peak viremia and day of onset of fever were compared between high and low exposure groups by Wilcoxon Rank Sum test. The Pearson’s correlation coefficient of morning T_r_ and log_10_ TCID_50_/ml was calculated. The duration of viremia and fever were not compared between groups, because animals in the high exposure group were euthanized on 4 DPF when viremia and fever were still present.

#### The relationship between the number of mosquito bites and infection outcomes

To examine if and how the probability of acquiring infection, *P*(inf), was affected by the number of infected mosquito bites received by a single host (*n*) in a short time frame, we fitted two probabilistic models to the experimental data, each representing a hypothesis of the nature of this possible dose-response relationship.

The models represent two distinct hypotheses:

i. P(inf) is independent of *n*:

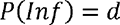
ii. P(inf) is dependent of *n*, where each subsequent infectious bite accrues the same per bite probability of infection (*b*), irrespective of earlier events:

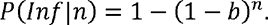

The first hypothesis can be regarded as the null hypothesis and denotes that the probability of infection is stable and not affected by the number of bites. It is here denoted by *d,* which can have any value between 0 and 100%. This hypothesis may be true especially for highly contagious pathogens where the received pathogen dose is not a limiting factor for transmission success. The second hypothesis reflects what is commonly adopted in mathematical models of mosquito-borne pathogens (Reiner et al., 2013). Herein, the probability for a single bite by an infectious mosquito to cause infection in a host, the vector-to-host transmission efficiency (*b*), is estimated. It assumes that, within an individual sheep, the probability of acquiring infection follows a Poisson process. Models *i* and *ii* were fitted to the experimental data by Maximum Likelihood Estimation and compared using AIC-scores (Burnham and Anderson, 2004).

The number of infected mosquitoes a lamb was exposed to (*n*) was derived differently between exposure groups. In the low exposure group, all mosquitoes were tested for infection and thus provide a direct measure of *n* (PCR positive bodies were used for the reported *b* calculation, because saliva results were not available for all mosquitoes). In the high exposure group, the number of infectious bites was estimated indirectly from a random sample of six mosquitoes per lamb. The number of engorged mosquitoes per individual in the high exposure group was multiplied with the proportion saliva positive mosquitoes to estimate the number of infectious bites (*n*) per lamb. A sensitivity analysis was performed to assess the impact of uncertainty in the estimate of the number of infectious mosquitoes (Suppl. Table 1).

#### Illustration of epidemiological implications

To explore if and under which circumstances RVFV transmission by mosquitoes with similar life history and transmission traits as *Ae. aegypti* could result in RVFV outbreaks, we applied the experimental estimates on the probability of transmission from mosquito to sheep (*b*) and sheep to mosquito (*c*) in a Ross-MacDonald-like framework for the basic reproduction number (Smith et al., 2007). The basic reproduction number here is defined as the average number of new infected sheep that are expected to arise from a single infected sheep over the course of its infectious period, in a fully susceptible population. It is a commonly used metric of transmission potential, as only if R_0_ surpasses 1, outbreaks may occur. It follows

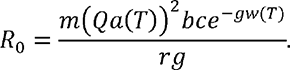

This definition follows from the several ‘hurdles’ a mosquito-borne pathogen needs to overcome to fulfil its transmission cycle (Figure 1). Parameter definitions and default values are depicted in Table 1. Here, the length of the extrinsic incubation period (*w*) and biting rate (*a*) are considered to change as a function of temperature. The biting rate is the inverse of the gonotrophic cycle (*gc*) length under the assumption that mosquitoes take a single bite per *gc*. Departure from this assumption is allowed for by including double feeding behavior (1 + *df*)⁄*gc*, where *df* denotes the proportion of mosquitoes that take a second bite during their *gc*. Mosquitoes are expected to remain infected for the remainder of their lives. Bites are assumed to be homogeneously distributed (i.e., every competent host has the same probability of getting bitten. The vector-to-host ratio (*m*) and the proportion of bites to be taken on a competent host (*Q*) are considered context specific, for example high vector-to-host ratios could be expected during rainy seasons, and bites on competent host would be high in a farm setting versus a city. We therefore explored the value of R_0_ in different host biting scenarios and with different vector-to-host ratios, while assuming a temperature equal to our experimental set-up (28°C). For illustrative purposes, we present two ‘extreme’ biting scenarios: vector opportunistically feeding on competent hosts (*Q* = 0.8) or on a preferred non-competent host (*Q* = 0.2). In this illustrative example, we assume all competent hosts to have transmission parameters similar to that of sheep. We illustrated the sensitivity of R_0_ to *b* by considering values of *b* from 0.05 to 0.95.

**Table 1:**
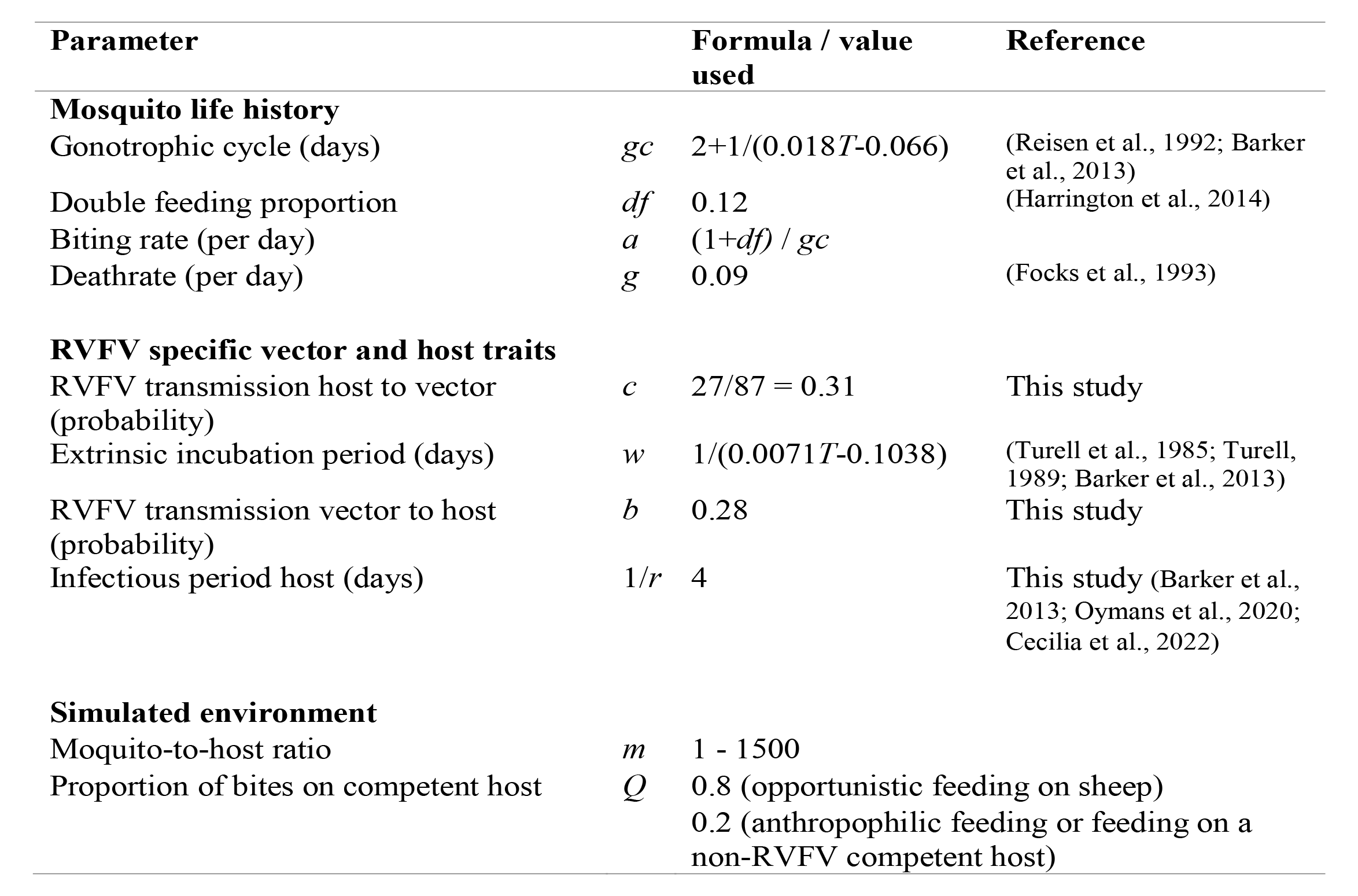
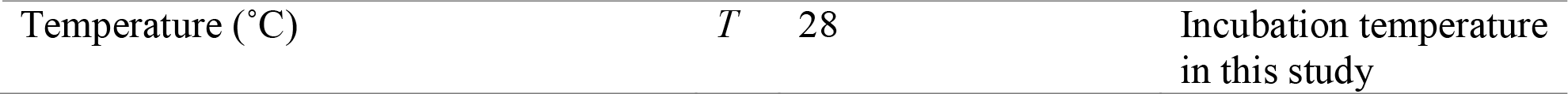
Description of parameters used in R_0_ calculation.

#### 4.4 Software

All animal observations were logged in an iVention notebook. Analyses were conducted with R Statistical Computing Software: *bbmle* (version: 1.0.24) was used for maximum likelihood estimation and AIC comparison, *binom* (version 1.1-1) for binomial confidence estimates calculation, and *ggplot2* (version 3.3.5) for visualizing data and model results (Dorai-Raj, 2014; Wickham, 2016, 2; Bolker and R Development Core Team, 2021; R Core Team, 2021).

#### 4.5 Data availability

R code for assessing the relationships between mosquito bites and infection is available at Open Science Framework https://osf.io/8vuax/ (DOI to follow).

## Results

### Host-to-mosquito transmission: obtaining RVFV exposed mosquitoes

To obtain infectious mosquitoes, approximately 1,000 female naive mosquitoes were allowed to take a blood meal from lambs that were needle inoculated with 10^5^ TCID_50_ RVFV two days prior. All five donor lambs had detectable fever at the time of mosquito feeding. A total of 641 mosquitoes fed on the lambs, these mosquitoes were pooled and maintained in an incubator until further use. Retrospectively, four out of five lambs had developed high viremia; one animal was viral RNA positive but had an RVFV titer at the limit of detection (Suppl. Table 2). The average RVFV titer in the blood of the lambs was 6.59 log_10_ TCID_50_/ml (range: 1.55, 7.15).

To estimate the proportion of mosquitoes that acquired infection (RVFV-positive bodies) and became infectious (RVFV-positive saliva), a subset of 30 mosquitoes was randomly selected for RT-qPCR testing on day 7 post initial feeding (experimental day −5). Due to logistic reasons, testing at a later time-point was not possible. Viral RNA was detected in 9 out of the 30 bodies tested, suggesting a 30% infection rate (95% confidence interval [95%CI]: 15 - 49%, Suppl. Table 3). Virus was not detected in corresponding saliva samples (0/30). Of note, the expected average extrinsic incubation period (*w*) at a 28°C incubation temperature is longer than 7 days; 10.5 days (Table 1) (Barker et al., 2013). Hence, we assumed that following another 5 days incubation (experimental day 0), virus would have reached the salivary glands in infected mosquitoes and one third of mosquitoes were expected to be infectious at day 0.

### Mosquito-to-host transmission

#### Exposure of naive lambs to a high or low number of infected mosquitoes

Based on the expected 30% infection ratio of the pool of RVFV-exposed mosquitoes and the aim to expose lambs of the low-exposure group to one bite containing RVFV on average, three mosquitoes were placed on each lamb. A retrospective analysis of these mosquitoes revealed that 9 out of 12 lambs were indeed exposed to a single bite by a mosquito with RVFV-positive bodies, whereas two received two and one received three bites (Figure 3, Suppl. Table 4). Of note, of the 9 lambs exposed to a single infected mosquito, 5 mosquitoes had detectable viral particles in their saliva, 3 did not and for 1 mosquito this information was not available.

**Figure 3:**
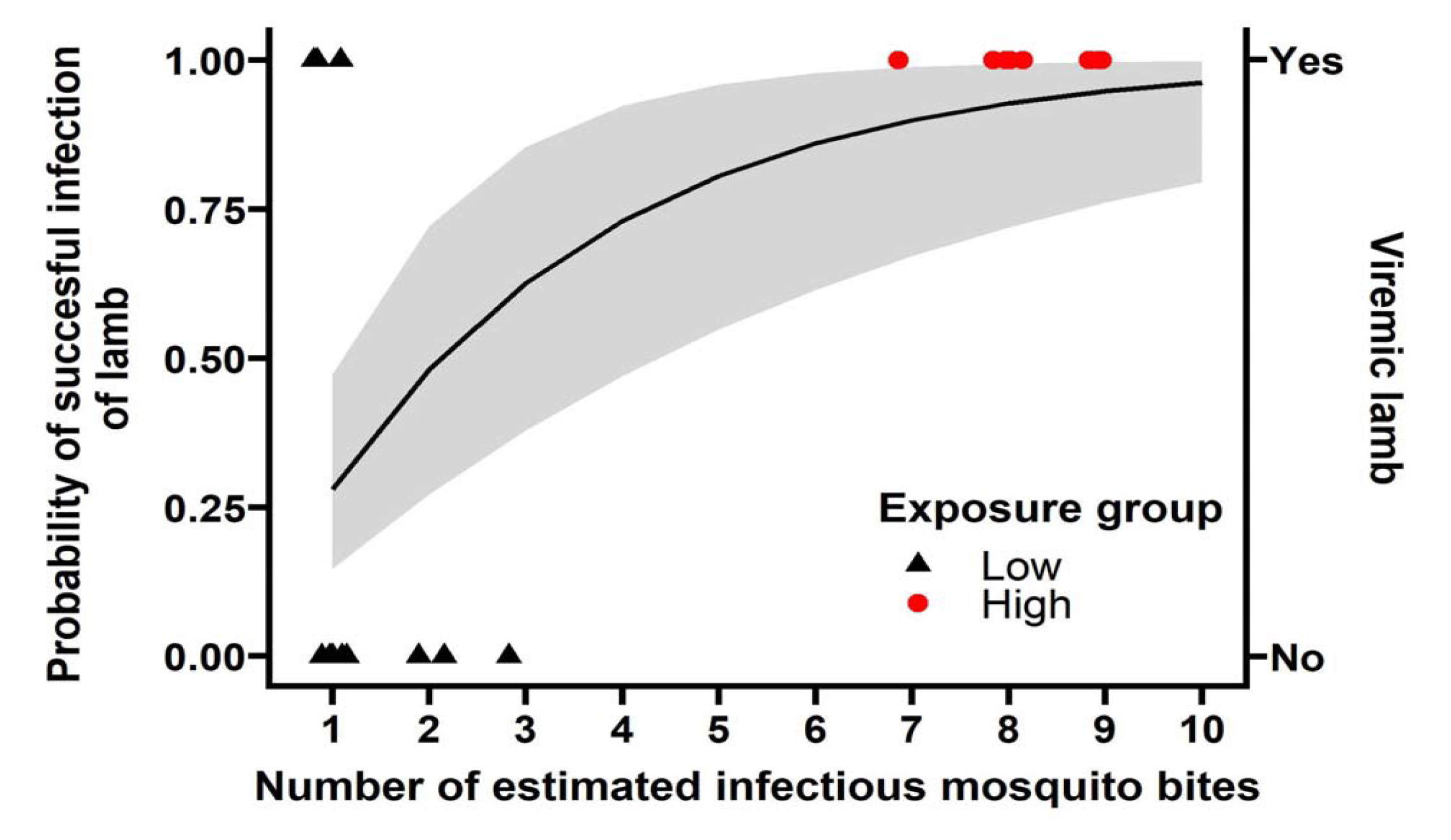
The number of bites matters for successful infection. Shapes represent the data collected (Suppl. Table 4), 9 individuals in the high exposure group (red) and 12 in the low exposure group (black) became viremic or not (y-axis on the right). The *b* estimate from the ‘bites matter’ model 0.280 (95%CI: 0.147, 0.473) was used to calculate the probability of infection per number of mosquito bites (black line, and grey ribbon represents its 95% confidence interval, y-axis on the left). Bites in the low exposure group represent PCR positive bodies, bites in the high exposure group represent the number of engorged mosquitoes expected to have RVFV in saliva.

In the high exposure group, lambs were exposed to 28-31 RVFV-exposed mosquitoes and 22 to 30 mosquitoes were engorged (Table 2, Figure 2). To estimate the number of infectious bites, saliva was collected from 33 of the blood fed mosquitoes from the low exposure group and a subset of the high exposure group (N=6 per lamb) immediately after feeding for a second time. Saliva from 27 of 87 mosquitoes had a positive TCID_50_ (31%, 95% CI: 22 - 42%). Taking the number of engorged mosquitoes and the proportion of RVFV positive saliva samples into account, lambs of the high exposure group were estimated to be exposed to 7-9 infectious bites (Table 2).

**Table 2:**
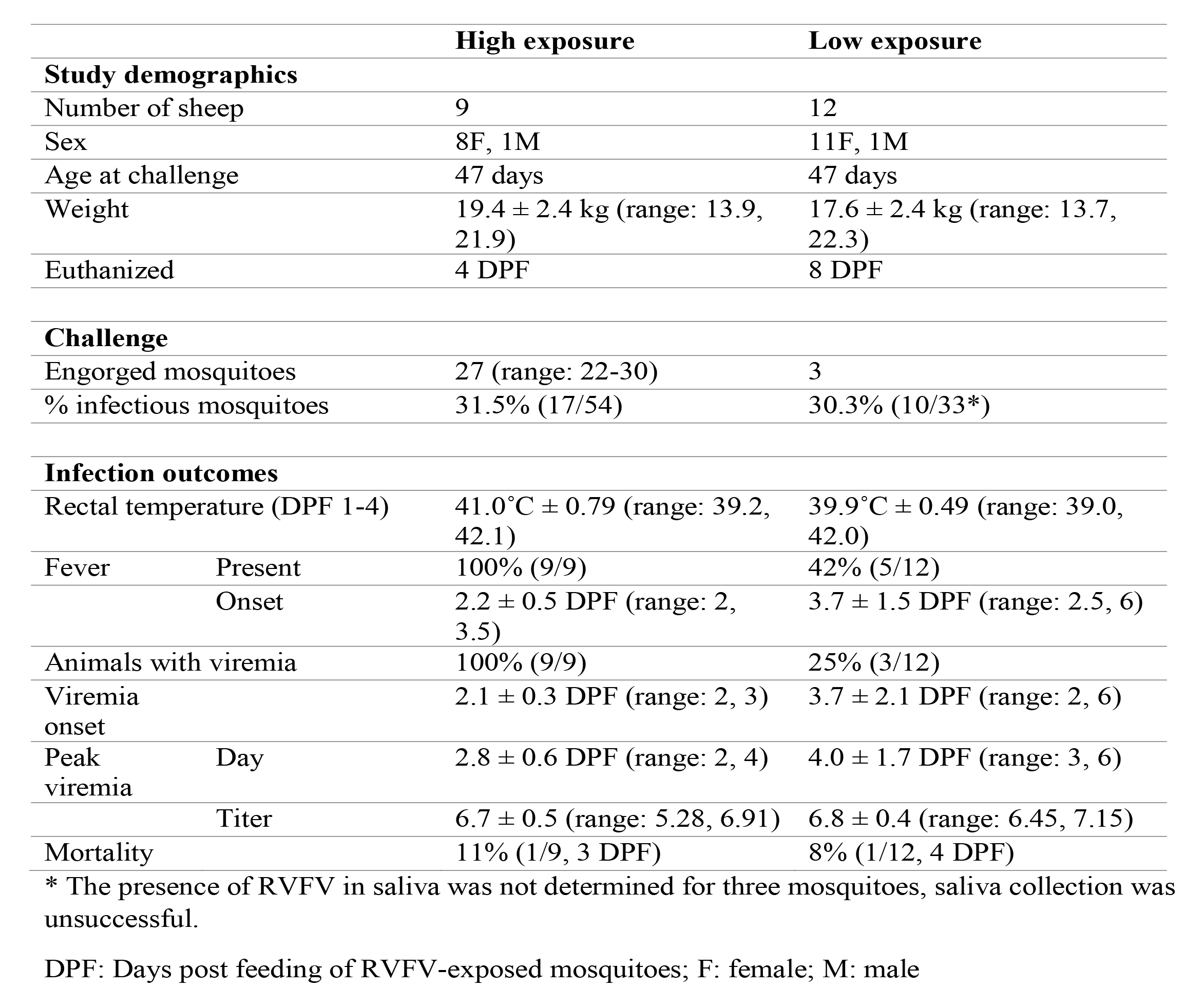
Summary of lamb characteristics, challenge and RVFV infection outcomes of high and low mosquito exposure groups. Mean ± SD are shown when applicable.

#### Clinical outcomes

##### High exposure group

Following mosquito challenge, all (9) high exposure animals presented with increased rectal temperatures from day 2 or 3 onwards (Fig. 4A). Furthermore, these animals presented with reduced feed intake and became lethargic. One animal succumbed to the infection and the other animals were euthanized at 4 DPF as planned. PCR on plasma samples, followed by virus isolation demonstrated that 9 out of 9 animals developed viremia with 8 animals presenting with a characteristic RVFV viremia curve starting at 2 DPF and 1 animal with a one day delay (Fig.4 C). In addition, viral RNA (Fig. 4G) and infectious virus were detected in all liver and spleen samples at the time of necropsy, except for the liver sample of animal #4620 in which no infectious virus was detected.

**Figure 4:**
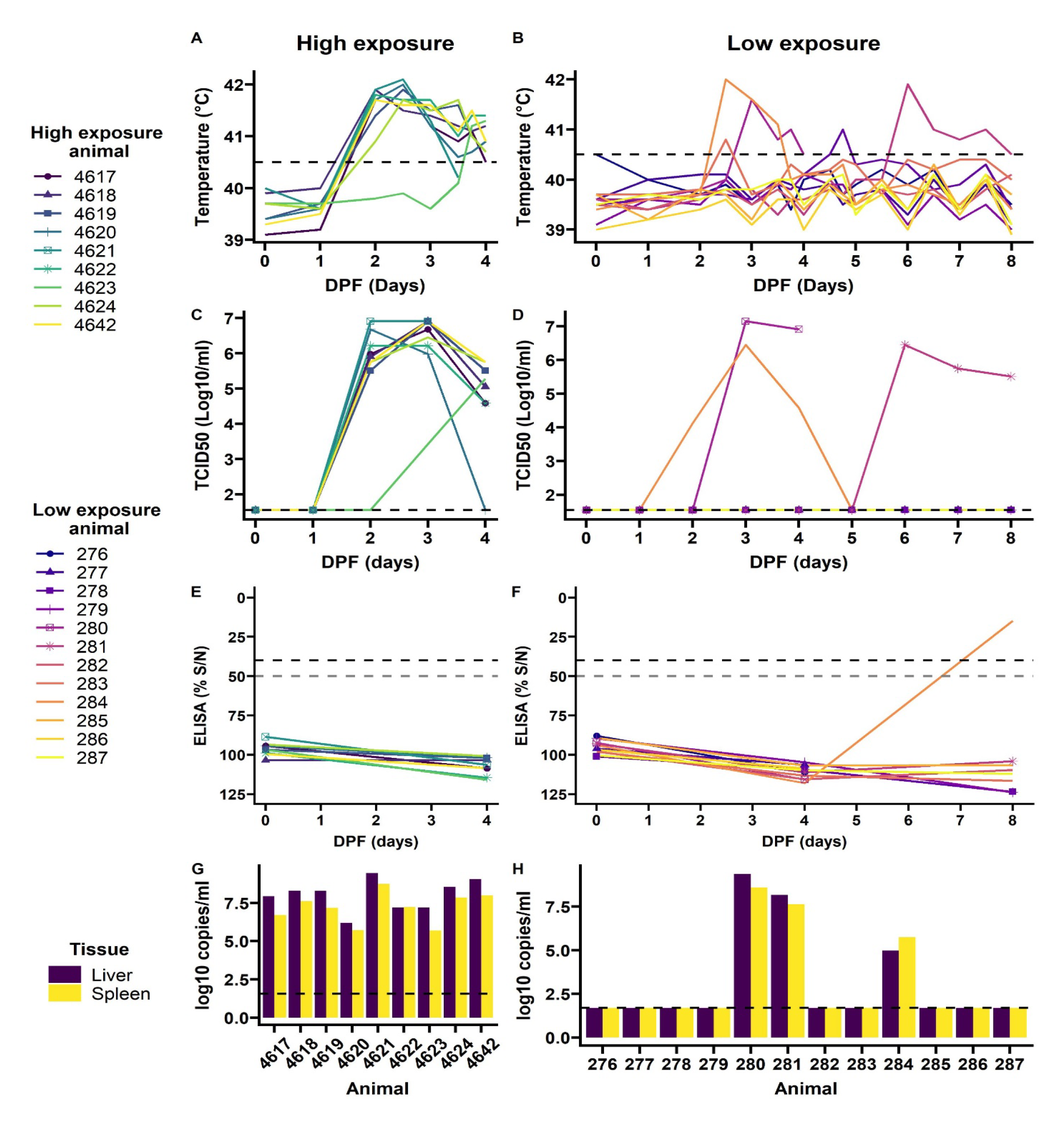
Outcomes of exposure to RVFV-exposed mosquitoes in individual lamb in high exposure (left) and low exposure (right) groups. Rectal temperatures were taken twice daily, and more frequent during 3 and 4 DPF (A and B). Temperatures over 40.5 C were considered a fever (dashed line). Blood samples were daily assessed for viremia development (C and D), limit of detection is shown with the dashed line. At day 0, 4 and 8 DPF anti RVFV antibody development was assessed by ELISA (E and F). A 50% signal was probable positive and 40% or less was considered positive (dashed lines) Upon euthanasia, RVFV presence was determined in 10% spleen and liver homogenates (G and H). PCR limit of detection shown in dashed line. Animal 4621 and 280 succumbed to infection on 3 and 4 DPF, respectively.

##### Low exposure group

In the low exposure group 9 out of 12 lambs did not present with any clinical signs of disease, nor presented with viremia and/or virus in liver and spleen samples at the time of necropsy, 8 DPF (Fig. 4B and 4D). However, 2 animals (#280 and #284) did present with characteristic features of RVFV infection starting at 2 DPF, which included fever, reduced feed intake and lethargy. Animal #280 succumbed to the acute infection at 4 DPF. A third animal (#281) presented with signs of disease later at 5 DPF. In line with the clinical signs all three animals presented with high viremia within one or two days post onset of symptoms. Furthermore, liver and spleen samples at the time of necropsy were highly positive for animal #280 and #281. Animal #284 had developed antibodies against RVFV-NP and cleared the virus (Fig. 4F and 4H).

#### Comparing natural infection between high and low exposure groups

After successful exposure to RVFV through mosquito bite(s) more animals in the high exposure group developed viremia compared to the low exposure group, 100% versus 25% respectively (FET *p*<0.001, Table 2, Figure 4). No difference in mortality was detected; one individual succumbed to RVF in each group (animal 4621 and 280, FET *p*=1, Table 2). All individuals that developed viremia also developed a fever (correlation coefficient 0.87, 95%CI: 0.82 - 0.90, t(146)=21.8, p<0.001). In the low exposure group, two additional animals had an afternoon or evening with fever 2 DPF and 4 DPF, but they did not have detectable viral RNA or virus in blood. No difference in peak viremia level was detected between the groups, peak viremias were 6.91 in the high exposure group and 7.15 log_10_ TCID_50_/ml in the low exposure group (t(2.31)=0.36, *p*=0.75). In the low exposure group the peaks appeared more variable, but no significant difference was detected in peak day between the two groups (W=21.5, *p*=0.07). None of the sheep had detectable anti-RVFV antibodies at the start of the study (Fig. 4E and 4F). The animals in the high exposure group were euthanized before antibodies could develop. No comparison was made with the low exposure group, where one individual had detectable antibodies 8 DPF.

#### Mosquito-to-host transmission efficiency (*b*)

Using the clinical outcomes and the individual-level information on number of infectious bites (Figure 3, Suppl. Table 4), we estimated the mosquito-to-host transmission efficiency (*b*). We tested whether the probability of acquiring infection is affected by the number of bites received, distinguishing two hypotheses: *i*) no impact of the number of bites on the acquisition of infection (the null hypothesis) or *ii*) the number of bites matters and each additional bite independently increases the infection probability. The model assuming bites to be independent of each other was best supported by the data (Model *ii*, AIC 19.4, delta AIC 11.3).

With model *ii*, we assume the infection probability to follow from a Poisson process in which each infectious bite is considered an independent attempt. Each independent attempt is estimated to result in infection with a probability (*b*) 0.280 (95%CI: 0.147, 0.473) (Figure 3). Using this model, we estimated that a host being exposed to two infectious bites has a near 50% chance of acquiring an infection (48.1%, 95%CI: 27.2, 72.2%).

For the reported estimate of *b* we used the number of body positive mosquitoes when referring to infectious mosquito bites in the low exposure group. For the high exposure group, we used a conventional approach: proportion of saliva positive mosquitoes of mosquitoes that were engorged. The number of mosquitoes with RVFV positive bodies was used in the low exposure group in order to deal with the missing datapoints for RVFV in saliva and the observation that one of the sheep exposed to mosquitoes without detectable RVFV in saliva became viremic (animal 281). Suggesting RVFV may have been present in saliva below the limit of detection.

Both model selection and estimates for *b* were robust for the use of different definitions of infections (TCID_50_ and PCR) and feeding status (exposed and engorged mosquitoes, Suppl. Table 1). Estimates for *b* ranged from 0.216 to 0.319 using different definitions of infectious bites (see Suppl. Table 1).

#### Illustration of epidemiological implications

Given estimated the mosquito-to-host transmission efficiency (*b*) based on our experimental studies we subsequently modelled under which circumstances outbreaks with *Ae. aegypti* involvement or alternative vectors with similar transmission and life history traits could occur. Specifically, we calculated the reproduction numbers (R_0_) for different mosquito-to-host densities and host-feeding preferences. We show that, for *Ae. aegypti* to be able to cause an outbreak in sheep, it needs to portray opportunistic biting behavior, i.e., in settings where sheep density is high compared to humans, it would need to feed proportionally more on sheep (or other competent ruminant species), despite its preference for feeding on humans (Figure 5). In such a scenario, high numbers of mosquitoes (mosquito-to-host ratio, *m* >200) are needed, even at 28°C, to surpass R_0_ =1. If 80% of bites were on humans (or other dead-end host for RVFV), higher numbers of *Ae. aegypti* would be needed for R_0_ to reach the critical value of unity (*m* >1000, Figure 5).

**Figure 5:**
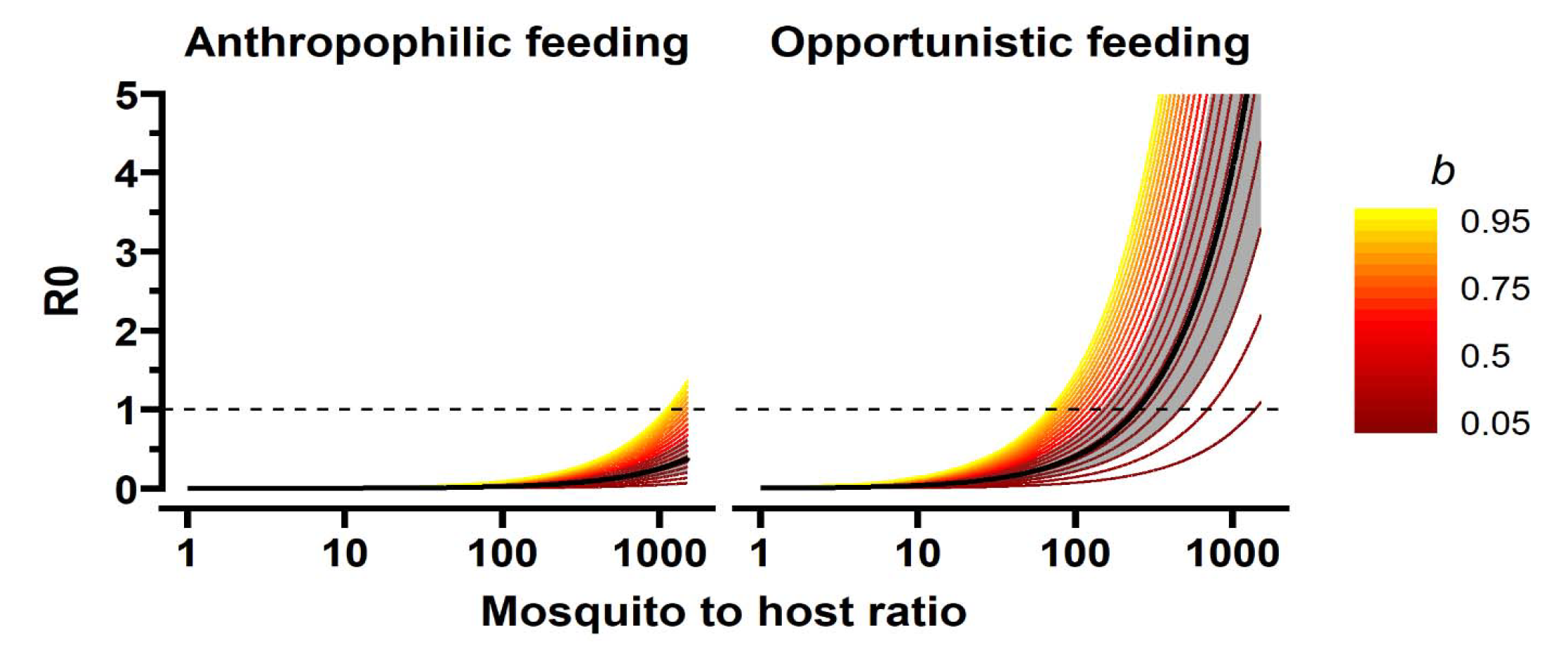
R_0_ for mosquito-to-host transmission efficiencies (*b)* in different contexts. The contexts represent: a mosquito to host ratio (*m*) from 1 to 1500 mosquitoes per lamb, and anthropophilic and opportunistic mosquito feeding behavior (*Q* = 20% versus 80% of bites on sheep). The bold black line represents the estimate derived from our experiment (Figure 2) and the grey shaded area includes the 95% confidence interval. The colored lines represent *b* from 0.05 to 0.95 from dark red to yellow. The vertical line represents the endemic equilibrium (R_0_ =1), values above this line indicate that an RVFV outbreak would occur in a naive population under the modeled scenarios whereby temperature was set at 28°C leading to an extrinsic incubation period (*w*) of 10.5 days and biting rate (*a*) of 0.26 bites per mosquito per day, mosquito death rate was 0.09 (an average lifespan of 11 days), the infectious period of lamb was 4 days and the host-to-vector transmission efficiency 0.31 (*c*).

## Discussion

Here, we performed a host-mosquito-host transmission experiment with RVFV in a controlled setting in which lambs were exposed to different numbers of infectious mosquitoes. We estimated the vector-to-host transmission efficiency of RVFV between Texel-Swifter lamb and *Ae. aegypti* mosquitoes. This work showed that the number of mosquito bites is a major factor in determining whether an animal develops an RVFV infection; all individuals exposed to 7+ infectious mosquito bites became infected, developed viremia and fever, whereas only a quarter of animals exposed to a single infectious mosquito on average was infected. The transmission efficiency from vector-to-host was estimated to be 28% (95% CI: 15 - 47%) and from host-to-vector 31% at peak viremia (95% CI: 22 – 42%). This study marks a unique empirical estimate of transmission efficiency from mosquitoes to a natural host species by a single mosquito bite, an important component of the transmission cycle of RVFV.

RVFV vector-to-host transmission efficiencies as determined under experimental conditions (*b*) can vary depending on the species and biotypes of mosquitoes, virus strain, routes of mosquito infection (e.g., membrane feeding, injection skipping the midgut-barrier, feeding on non-natural host species), and host species used (Turell and Bailey, 1987; Turell et al., 2008). Here we used the proportion of Texel-Swifter lambs that became viremic as our primary outcome measure for successful transmission. With 3 out of 12 sheep infected after exposure to a single bite by RVFV positive mosquitoes, the transmission of RVFV from *Ae. aegypti* in this study was higher than that of other studies using *Ae. aegypti* as a vector. For instance, 1 out of 7 hamsters (14%) succumbed to RVFV after a bite from *Ae. aegypti* with disseminated infection (Turell et al., 2008). The transmission efficiency is likely reflected by this proportion, as hamsters experience 100% mortality to infection. Further, in early host-vector-host transmission experiments, no transmission was observed from 8 exposed *Ae. aegypti* mosquitoes (5 of which contained RVFV) to mice and lamb (Smithburn et al., 1949). It was however not reported if individual or small groups of mosquitoes were used for animal exposure. Such intricacies make the estimate of *b* difficult to compare to other studies. For example, 55, 84 and 100% transmission of RVFV was observed from African *Ae. palpalis*, *Culex antennatus* and *Cx. pipiens* with disseminated infection to hamsters, but this could have been based on 1 to up to 5 mosquitoes per animal (Turell et al., 2008). Furthermore, care should also be taken in extrapolating vector-to-host probability of transmission by a single bite (*b*) to other ruminant species or other sheep breeds, for instance those present in RVFV endemic countries. Estimates for other species are expected to be lower as sheep are deemed most susceptible to RVFV infection. In addition, we did not consider probing (i.e., unsuccessful biting attempts) as an exposure event, even though these could potentially also contribute to transmission (Matsuoka et al., 2002; Styer et al., 2007). We also did not consider body positive mosquitoes for our denominator in the high exposure group. Instead we used saliva positive mosquitoes. This may result in an overestimation of *b*, because too few mosquitoes are considered capable of transmitting virus (*n*). The method of detecting virus in saliva has imperfect sensitivity (Gloria-Soria et al., 2022). This was also demonstrated by the animal in the low exposure group that did become viremic after exposure to mosquitoes without detectable viral particles in saliva. Using more liberal infection rates for mosquitoes, i.e., using the number of mosquitoes with PCR positive bodies, results in somewhat lower levels of *b* (24%, 95%CI: 12- 44%, Suppl. Table 1).

This study re-enforces two topics of mosquito-borne pathogen transmission at the level of the individual host: i) the number of (infectious) mosquito bites individual animals receive determine the probability of acquiring infection, ii) within hosts, if infection is successful and viremia develops, viremia peaks appear similar between low and high exposed individuals. Development of RVF and RVFV trajectories in the high exposure group were similar to Wichgers Schreur et al.; rectal temperatures and RVFV titers were strongly correlated and both peaked on 2-3 DPF (Wichgers Schreur et al., 2021). This relative short and intense viremia is seen in several other arboviral infections (Binn et al., 1967; Oesterle et al., 2009). The phenomena is often interpreted in light of the virus’ tradeoff between magnitude and duration of viremia. This tradeoff has been observed in experiments with varying dosages, where high dosages result in high-peak, short-lasting viremia, whereas low dosages result in longer lasting viremias that peaks at lower levels. The mechanism behind this dose response relationship is not completely clear and may well vary between pathogens. One explanation is that high initial viremia accelerates both initial virus replication and the recruitment of immune cells. While observed in many arboviruses, the evidence for such a relationship in arboviruses is not consistent (Althouse and Hanley, 2015). One counter example, for instance, follows from West Nile virus studies, which found no difference in peak viral titers between high and low dose groups (Oesterle et al., 2009; VanDalen et al., 2013). Similar to these observations in West Nile virus viral kinetics, we found no indication that peak titers were affected by the level of mosquito exposure. Two interesting observations in the low exposure group could be examined further in future studies: the seemingly later onset of viremia, as well as the slow decline in viremia in one animal in this group. The later onset of viremia may be caused by initial local (skin and or lymph node) replication, before the virus reached the liver and initiated full viremia. Notably, the heterogeneity observed within the low exposure group may in part result from differences in viral dosage in mosquito saliva. Due to the natural exposure route used in our study we were not able to control exposure dose. The possible variation in exposure dose could have blurred part of the dose response relationships, if present. In general, gaining understanding of the range of infection outcomes that may arise from mosquito-borne RVFV infection and how this is coupled to heterogeneity within mosquitoes (i.e., viral dose) can help decipher the transmission landscape of this virus.

We estimated an upper boundary for the host-to-mosquito transmission efficiency (*c*) by estimating this parameter at peak viremia of needle-inoculated sheep (31%, 95%CI: 22 - 42%). Peak viremia is likely the moment hosts are most contagious. A host’s transmission potential over their full infectious period (*c*r^-1^,* net infectiousness) will be lower than expected based on this estimate of *c*. Estimates of net infectiousness derive from the full trajectory of increase and decline of infectious viral particles over the course of the infectious period. The viremia levels are coupled with estimations of how these levels relate to transmission efficiency. This dose response relationship has been estimated for RVFV based on a meta-analysis of available literature and presents a composite estimate across vertebrate species and the *Aedes* genera (Cecilia et al., 2022). Our results (31% from 4.35 to 7.15 log10 TCID50/ml) are consistent with the described relationships. Estimates of *c* across the full infectious period would aid in getting a better assessment of the net infectiousness and associated dose response relationships. In this study only one full viral trajectory was observed for the low exposure group. The absence of additional complete viral trajectories limits the ability to estimate net infectiousness (using published dose response relationships) and assess how mosquito exposure affects transmission potential.

Using the transmission efficiency estimates from our host-mosquito-host model system in a RossMcDonald-type R_0_ calculation, illustrated that RVFV (vectored by a mosquito with similar transmission potential as *Ae. aegypti*) could invade naive areas if favorable conditions are met. These conditions include temperatures conducive to efficient transmission, sufficiently high mosquito-to-host ratios, and large proportions of mosquito bites on competent hosts. The local vulnerability to sustained pathogen transmission is particularly sensitive to the vector-to-host ratio. This is because a sheep needs to be bitten twice to fulfil the transmission cycle: once to become infected and once to contribute to onward transmission of the pathogen. In that transmission chain the number of bites received by a sheep is proportional to the vector-to-host ratio (i.e,, while a mosquito will not feed more often when more hosts are available, the chance for an individual sheep to be fed on is higher if more there are sizably more mosquitoes than sheep). All these conditions are context specific. For example, most mosquitoes have a host preference, but many will divert from this preferred species depending on local host availability. This was illustrated in (Tandon, Neelam and Ray, Sudipta, 2000), where 82% of highly anthropophilic *Ae. aegypti* caught in a cattle-shed had fed on cattle. Furthermore, here we used a theoretical model whereby bites were homogeneously distributed over a host species, i.e., every individual receives the same number of bites. In nature bites are often heterogeneous distributed, with some individuals receiving many bites and others few, affecting pathogen invasion and transmission at population levels (Woolhouse et al., 1997). Together this reinforces that field observations are essential for translation of experimental data to epidemiological parameters and outcomes.

With this host-mosquito-host transmission model we aimed to reproduce the RVFV transmission cycle in a way that best mimics natural virus exposure. Mirroring natural transmission cycles in studies create challenging logistics; host-mosquito-host studies require dedicated facilities, a relatively long timeline, two carefully timed mosquito feeding events, the survival of mosquitoes to feed a second time, among other challenges. To enhance feasibility, deviations from natural systems must be made in model systems. For example, the sheep used in our study are native to the Netherlands and not common in RVFV endemic countries. Further, although this lab-reared *Ae. aegypti* strain can transmit RVFV, other *Aedes* species are more important for RVFV transmission in endemic areas (Turrell et al., 2008. We chose *Aedes aegypti* as our model species, because this species is easy to maintain in laboratory settings, willing to feed at least twice in captivity and capable of transmitting RVFV to ruminant species (Wichgers Schreur et al., 2021, and this paper). Replacing host and vector species with the species of interest could improve transmission estimates. However, with a relative promiscuous virus, like RVFV, it is impossible to test all possible, and even all relevant, combinations. Moreover, many mosquito species are not suitable for lab rearing and/or are not capable of surviving in laboratory settings to live past the extrinsic incubation period. Furthermore, we tested at 28°C, higher and lower temperatures likely affect transmission efficiency estimates as well (Tesla et al., 2018). This temperature – transmission relationship could be further explored in future studies. Furthermore, the extended observation period in the low exposure group highlighted that, despite animal welfare and cost concern, prolonged observations may be warranted to address questions on the progression of infection after low exposure. Specifically, the relationship between exposure dose, viremia (including how viremia levels decline) and onward transmission can be extrapolated more precisely. Thus, although model systems cannot capture reality, they are useful to explore coupled heterogeneities (exposure, viremia, transmission) and establish parameter estimates that can be used in epidemiological models.

Mathematical model parameters are often informed by experiments not designed for this purpose. Consequently, estimates may misrepresent the efficiency of transmission or outcomes of infections, for instance due to the use of high inoculation dosages or artificial transmission routes. In this experimental sheep-mosquito-sheep design, we have successfully reproduced RVFV infection in sheep by the bite of a single mosquito, the most likely route ruminants become exposed to RVFV. This design could be used for future research on different host and vector combinations and expanded to examine the drivers behind the heterogeneous outcomes, such as observed in the low exposure group. Individual-level transmission parameter, such as described here, should always be put in the ecological context where the virus is present or at risk of emerging. Teams of epidemiologists, entomologists, ecologists, and virologists, among others, can help to synthesize knowledge across disciplines and put these in an epidemiological perspective meaningful for animal and public health decision makers.

### Author contributions

Conceptualization: QB, JK, MCMJ, CJMK, LK, PJWS; Data Curation: PJWS, QB, GMB; Formal Analysis: QB, GMB; Funding Acquisition: JK, QB, GMB, MCMJ; Investigation: JK, PJWS, LK, RPMV; Visualization: QB, GMB; Writing – Original Draft preparation: QB, GMB, PJWS; Writing – Review & Editing: JK, CJMK, LK, RPMV, MCMJ

## Acknowledgements

We acknowledge the privilege that we were able to conduct this study with animals. We are thankful to Jet Kant, Sandra van de Water and animal care staff for technical assistance respectively caring for the animals. We thank Tessa Visser and Pieter Rouweler for their help in rearing *Aedes aegypti*.

