## Supplemental files for "Quantifying Rift Valley fever virus transmission efficiency in a lamb-mosquito-lamb model"

### Supplemental Table 1: Estimate of *b* under different mosquito infection criteria

Bold and yellow is reported in the manuscript.

| Hypothesis | *b* | CI_low | CI_high | AIC | Low_exposure | High_Exposure_type | High_Exposure_proportion | High_Exposure_estimate |
|---|---|---|---|---|---|---|---|---|
| H1 | 0.29 | 0.14 | 0.54 | 15.82 | Set to 1 | Exposed | PCR_body | Mean |
| H1 | 0.29 | 0.14 | 0.54 | 15.86 | Set to 1 | Exposed | TCID50_saliva | UpperCI |
| H1 | 0.29 | 0.14 | 0.54 | 15.92 | Set to 1 | Engorged | PCR_body | Mean |
| H1 | 0.30 | 0.15 | 0.55 | 16.02 | Set to 1 | Engorged | TCID50_saliva | UpperCI |
| H1 | 0.32 | 0.16 | 0.56 | 16.33 | Set to 1 | Exposed | TCID50_saliva | Mean |
| H1 | 0.18 | 0.08 | 0.33 | 16.35 | Set to 1 | Exposed | PCR_body | UpperCI |
| H1 | 0.33 | 0.17 | 0.56 | 16.54 | Set to 1 | Engorged | TCID50_saliva | Mean |
| H1 | 0.18 | 0.09 | 0.34 | 16.55 | Set to 1 | Engorged | PCR_body | UpperCI |
| H1 | 0.33 | 0.17 | 0.56 | 16.58 | Set to 1 | Exposed | PCR_body | LowerCI |
| H1 | 0.34 | 0.18 | 0.57 | 16.79 | Set to 1 | Engorged | PCR_body | LowerCI |
| H1 | 0.36 | 0.20 | 0.59 | 17.39 | Set to 1 | Exposed | TCID50_saliva | LowerCI |
| H1 | 0.37 | 0.20 | 0.60 | 17.68 | Set to 1 | Engorged | TCID50_saliva | LowerCI |
| H1 | 0.22 | 0.10 | 0.43 | 17.75 | PCR positive bodies | Exposed | PCR_body | UpperCI |
| H1 | 0.22 | 0.10 | 0.43 | 17.86 | PCR positive bodies | Engorged | PCR_body | UpperCI |
| H1 | 0.24 | 0.12 | 0.44 | 18.25 | PCR positive bodies | Exposed | PCR_body | Mean |
| H1 | 0.24 | 0.12 | 0.44 | 18.32 | PCR positive bodies | Exposed | TCID50_saliva | UpperCI |
| H1 | 0.25 | 0.12 | 0.45 | 18.42 | PCR positive bodies | Engorged | PCR_body | Mean |
| H1 | 0.25 | 0.13 | 0.45 | 18.60 | PCR positive bodies | Engorged | TCID50_saliva | UpperCI |
| H1 | 0.27 | 0.14 | 0.47 | 19.10 | PCR positive bodies | Exposed | TCID50_saliva | Mean |
| **H1** | **0.28** | **0.15** | **0.47** | **19.42** | **PCR positive bodies** | **Engorged** | **TCID50_saliva** | **Mean** |
| H1 | 0.28 | 0.15 | 0.48 | 19.48 | PCR positive bodies | Exposed | PCR_body | LowerCI |
| H1 | 0.29 | 0.15 | 0.48 | 19.79 | PCR positive bodies | Engorged | PCR_body | LowerCI |
| H1 | 0.31 | 0.17 | 0.50 | 20.66 | PCR positive bodies | Exposed | TCID50_saliva | LowerCI |
| H1 | 0.32 | 0.17 | 0.51 | 21.06 | PCR positive bodies | Engorged | TCID50_saliva | LowerCI |
| **H2** | **0.57** | **0.36** | **0.77** | **30.68** | | | | |

### Supplemental Table 2: Donor sheep Rift Valley fever virus titer

| Individual | $Log_{10}$ copies / ml | $Log_{10}$ $TCID^{50}$/ml | Mosquitoes fed |
|---|---|---|---|
| 271 | 8.07 | 4.35 | 288 |
| 272 | 9.13 | 5.98 | 102 |
| 273 | 8.70 | 7.15 | 162 |
| 274 | 9.77 | 6.45 | 41 |
| 275 | 3.63 | 1.55 | 48 |

### Supplemental Table 3: Pre-challenge mosquito infection check

To estimate RVFV infection (positive bodies) and dissemination (positive saliva) rates, 30 mosquitoes were randomly selected for RT-qPCR testing on day 7 post feeding (experimental day -5).

| **Mosquito** | **Body** | **Saliva** |
|---|---|---|
| 1 | Neg | Neg |
| 2 | Neg | Neg |
| 3 | Neg | Neg |
| 4 | Neg | Neg |
| 5 | 2.50E+05 | Neg |
| 6 | 1.37E+05 | Neg |
| 7 | Neg | Neg |
| 8 | Neg | Neg |
| 9 | Neg | Neg |
| 10 | Neg | Neg |
| 11 | 4.90E+04 | Neg |
| 12 | Neg | Neg |
| 13 | Neg | Neg |
| 14 | 7.52E+04 | Neg |
| 15 | Neg | Neg |
| 16 | Neg | Neg |
| 17 | Neg | Neg |
| 18 | Neg | Neg |
| 19 | Neg | Neg |
| 20 | Neg | Neg |
| 21 | 1.91E+05 | Neg |
| 22 | Neg | Neg |
| 23 | Neg | Neg |
| 24 | 2.50E+05 | Neg |
| 25 | 4.89E+04 | Neg |
| 26 | Neg | Neg |
| 27 | 7.20E+05 | Neg |
| 28 | Neg | Neg |
| 29 | Neg | Neg |
| 30 | 1.29E+02 | Neg |
| **Total** | **9/30** | **0/30** |

### Supplemental Table 4: Summary of infectious bites and outcome

The number of animals that became viremic after a certain number of infectious bites, and the total number of animals per exposure.

| **Group** | **Low exposure** | | | **High exposure** | | |
|---|---|---|---|---|---|---|
| **Number of infectious bites** | 1 | 2 | 3 | 7 | 8 | 9 |
| **Number of viremic / exposed animals** | 3 / 9 | 0 / 2 | 0 / 1 | 1 / 1 | 4 / 4 | 4 / 4 |

### Supplemental Figure 1

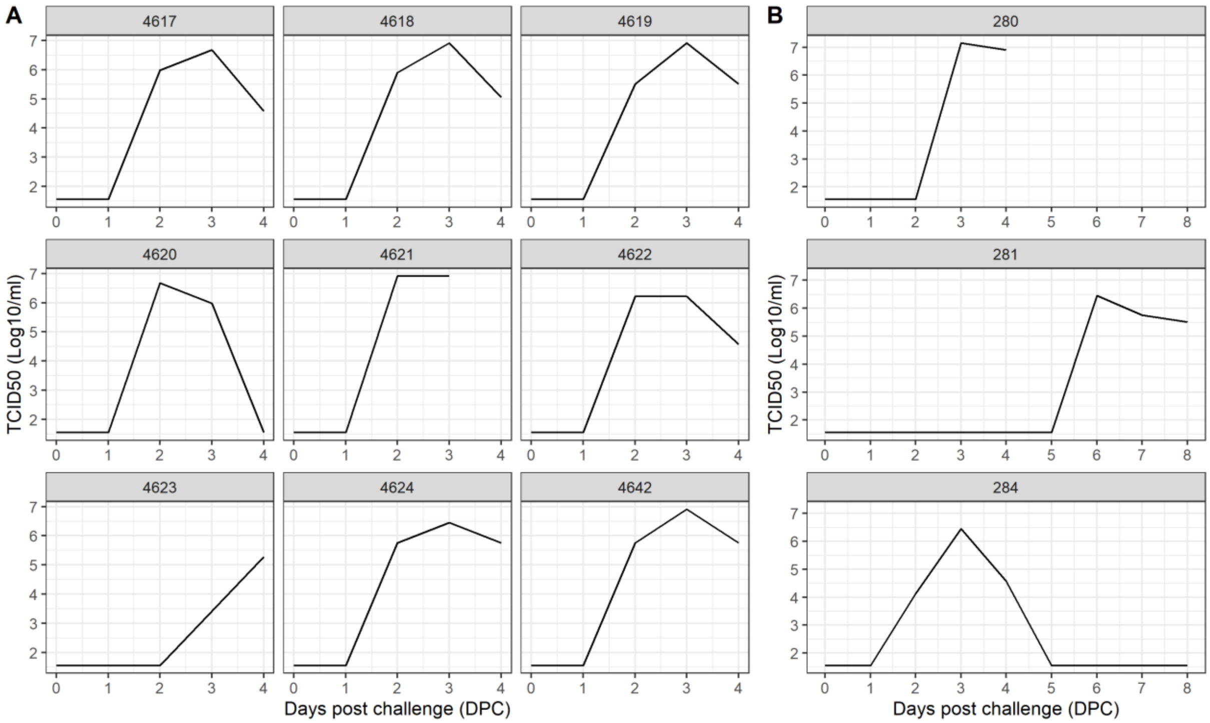

Rift Valley fever virus viremia trajectory of individual sheep in the high exposure (A) and low exposure (B) groups. Blood samples were taken daily. Animal 4621 and 280 succumbed to infection.
